# Targeting SCD triggers lipotoxicity of cancer cells and enhances anti-tumor immunity in breast cancer brain metastasis mouse models

**DOI:** 10.1101/2024.05.06.592766

**Authors:** Alessandro Sammarco, Giorgia Guerra, Katharina M. Eyme, Kelly Kennewick, Yu Qiao, Joelle El Hokayem, Kevin J. Williams, Baolong Su, Valentina Zappulli, Steven J. Bensinger, Christian E. Badr

## Abstract

Breast cancer brain metastases (BCBM) are a significant cause of mortality and are incurable. Thus, identifying BCBM targets that reduce morbidity and mortality is critical. BCBM upregulate Stearoyl-CoA Desaturase (SCD), an enzyme that catalyzes the synthesis of monounsaturated fatty acids, suggesting a potential metabolic vulnerability of BCBM. In this study, we tested the effect of a brain-penetrant clinical-stage inhibitor of SCD (SCDi), on breast cancer cells and mouse models of BCBM. Lipidomics, qPCR, and western blot were used to study the in vitro effects of SCDi. Single-cell RNA sequencing was used to explore the effects of SCDi on cancer and immune cells in a BCBM mouse model. Pharmacological inhibition of SCD markedly reshaped the lipidome of breast cancer cells and resulted in endoplasmic reticulum stress, DNA damage, loss of DNA damage repair, and cytotoxicity. Importantly, SCDi alone or combined with a PARP inhibitor prolonged the survival of BCBM-bearing mice. When tested in a syngeneic mouse model of BCBM, scRNAseq revealed that pharmacological inhibition of SCD enhanced antigen presentation by dendritic cells, was associated with a higher interferon signaling, increased the infiltration of cytotoxic T cells, and decreased the proportion of exhausted T cells and regulatory T cells in the tumor microenvironment (TME). Additionally, pharmacological inhibition of SCD decreased engagement of immunosuppressive pathways, including the PD-1:PD-L1/PD-L2 and PVR/TIGIT axes. These findings suggest that SCD inhibition could be an effective strategy to intrinsically reduce tumor growth and reprogram anti-tumor immunity in the brain microenvironment to treat BCBM.

## Introduction

Approximately 20% of cancer patients develop brain metastases (BM), the majority of which result from lung (20-56%), breast (5-20%), melanoma (7-16%), kidney (4-11%), and colorectal cancer (1-12%) (1–6). Therapies include combinations of surgical resection, radiotherapy, chemotherapy, and immunotherapy. Despite multimodal therapies, individuals with BM have a poor prognosis, with an overall 2-year survival below 10% (1,7). Two major factors contribute to the poor outcome of therapy (8). First, the molecular profile of BM could differ from that of the primary tumor (9). This would inevitably impact the metabolic profile and response to therapy of these BM. Second is the blood-brain barrier (BBB) which impedes the delivery of most drugs that would be effective for extracranial tumors (10–12).

The brain microenvironment forces metastatic cells to undergo lipid metabolic adaptations to survive and proliferate in the brain. Metabolic rewiring highlights potential vulnerabilities that can be exploited to develop targeted therapies. It has been shown that considerably lower levels of triacylglycerols (TAGs) are available in the brain than in the tissue environment of the primary tumor. As a result, metastatic cells must synthesize requisite lipids *de novo* to meet the anabolic demands of proliferation (13). In support of this idea, it has been shown that BM from human breast cancer (HBC) upregulate several enzymes involved in fatty acid metabolism, including Stearoyl-CoA Desaturase (SCD). This key enzyme converts saturated fatty acids (SFAs) into monounsaturated fatty acids (MUFAs) (12,13). The balance between SFAs and MUFAs contributes to the biophysical properties of cell membranes, such as fluidity, permeability, and the spatial assembly of membrane properties (14). SCD also regulates the pool of MUFAs, which serve as building blocks for more complex lipids (*e.g.*, phospholipids and TAGs) that are critical for cell survival (14). The heightened expression of SCD and an increased reliance on *de novo* synthesis of MUFAs in breast cancer brain metastases (BCBM) has led to the hypothesis that this metabolic enzyme could be an attractive therapeutic target in metastatic disease. However, SCD inhibitors (*e.g.*, CAY10566, A939572, and MF-438) have a limited capacity to cross the BBB, hampering their clinical relevance for BM. The development of the brain-penetrant SCD inhibitor, YTX-7739 (15) (hereafter referred to as SCDi), has warranted renewed interest in SCD as a target for brain malignancies. Recently, we showed that SCD is essential for glioblastoma (GBM) tumor growth (16), and demonstrated the therapeutic efficacy of SCDi in GBM mouse models (17).

Another important factor underlying the growth and survival of brain metastatic tumors is their interaction with the immune system. Several immune subsets exist within the brain tumor microenvironment (TME) that can have distinct and sometimes opposing roles. Increasing evidence suggests that fatty acid homeostasis in immune cells represents a key regulatory node to control inflammation, and manipulation of fatty acid metabolism has significant effects on immune cell functions (18–20). While the efficacy of pharmacological inhibition of SCD was already investigated in extracranial tumors and GBM (17,21,22), its effect on cancer cells and anti-tumor immunity in the BCBM TME has not been examined. Therefore, we studied the effect of a brain-penetrant clinical-grade SCD inhibitor in BCBM and examined the impact of this inhibitor at the single-cell level on the cancer and immune cells in the TME.

## Results

### SCD inhibition reshapes the lipid composition of breast cancer cells

To confirm the on-target effect and evaluate the influence of SCDi on fatty acid synthesis flux in our model, HBC cells were cultured in ^13^C-glucose-containing media. After 48h of labeling, lipids were extracted and analyzed by GC-MS to determine isotope enrichment into long-chain fatty acids. Isotopologues were then modeled using isotope spectral analysis (ISA) to assess the contribution of synthesis to the total cellular fatty acid pool in the cells. As expected, pharmacological inhibition of SCD led to a significant dose- and time-dependent increase of synthesized SFA 18:0, an SCD substrate, and a corresponding decrease in SCD products 16:1 and 18:1 (**Figures 1A, S1A-B**). Accordingly, the fatty acid desaturation index (FADI) was significantly reduced in cells treated with SCDi (**Figures 1A and S1B**). To determine the influence of pharmacological inhibition of SCD on the lipidome of HBC cells, we performed shotgun lipidomics on brain-tropic MDA-MB-231-BR (hereinafter referred to as MDA-BR) treated with SCDi. Approximately 900 individual lipid species from 17 lipid subclasses were measured. Cluster analysis of individual lipid species revealed that SCDi treatment markedly changed the lipid composition of MDA-BR cells (**Figures 1B, S1C-E**). Analysis of SFAs and MUFAs incorporation into phospholipids (PLs), triacylglycerides (TAGs), and cholesteryl esters (CEs) revealed that SCDi treatment showed an increased proportion of SFAs, and a corresponding decreased proportion of MUFAs, consistent with SCD inhibition (**Figures 1C and S1F**). Of note, the difference in SFAs between control and SCDi was much higher in TAGs (47% vs 73%, respectively) and in CEs (33% vs 57%, respectively), compared to PLs (38% vs 43%, respectively). Interestingly, in SCDi-treated cells, the percentage of polyunsaturated fatty acids (PUFAs) increased only in PLs, while decreased in TAGs and CEs (**Figure 1C**). These data confirm that SCDi inhibits SCD-dependent MUFA synthesis and can markedly reshapes the breast cancer cell lipidome.

**Figure 1.**
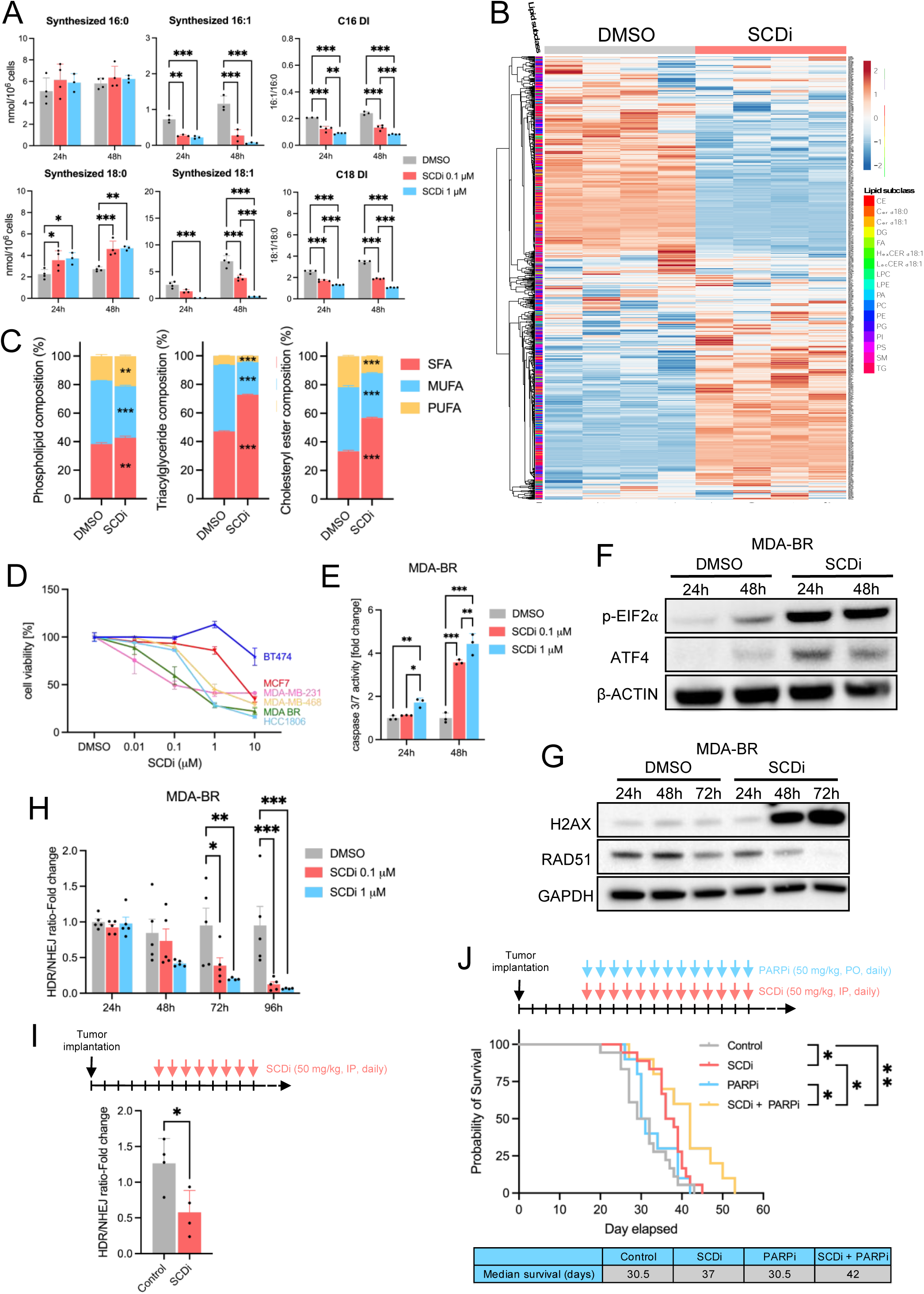
Inhibition of SCD induces ER stress, is associated with DNA damage, and prolongs survival in a mouse model of BCBM. (A) Gas chromatography/mass spectrometry of HCC1806 treated with SCDi for 24h and 48h, measuring synthesized palmitic acid (16:0), palmitoleic acid (16:1), stearic acid (18:0), and oleic acid (18:1). On the right, the desaturation index (DI - ratio 16:1/16:0 and 18:1/18:0) is shown. Mean ± SD (Two-way ANOVA, Tukey’s test). (B) Heat map of all individual lipids quantified using shotgun lipidomics of MDA-BR treated with SCDi (1μM) for 48h. CE, cholesteryl esters; Cer, ceramides; DG, diacylglycerols; FA, free fatty acids; HexCER, hexosylceramides; LacCER, lactosylceramides; LPC, lysophosphatidylcholines; LPE, lysophosphatidylethanolamines; PA, phosphatidic acid; PC, phosphatidylcholines; PE, phosphatidylethanolamines; PG, phosphatidylglycerols; PI, phosphatidylinositol; PS, phosphatidylserines; SM, sphingomyelin; TG, triacylglycerols. (C) Stacked histograms showing saturated fatty acids (SFAs), monounsaturated FAs (MUFAs), and polyunsaturated FAs (PUFAs) composition in phospholipids, triacyglicerides, and cholesteryl esters of MDA-BR treated with SCDi (1μM) for 48h, measured by shotgun lipidomics. Mean ± SD (Student’s t test). Statistics refers to DMSO vs SCDi. (D) Cell viability of breast cancer cells treated with SCDi for 72h. Mean ± SD. (E) Caspase 3/7 assay of MDA-BR treated with SCDi. Mean ± SD (Two-way ANOVA, Tukey’s test). (F-G) Western blot for (F) ER stress and (G) DNA damage markers in MDA-BR treated with SCDi (1μM). (H-I) Vargula luciferase (Vluc) and Gaussia luciferase (Gluc) assay measured as fold change in HDR/NHEJ in (H) MDA-BR treated with SCDi in vitro and (I) plasma of mice subcutaneously implanted with MDA-BR and treated with SCDi. (H) Mean ± SEM (Two-way ANOVA, Tukey’s test). (I) Mean ± SD (Student’s t test). (J) Overall survival of mice implanted with MDA-BR intracranially and treated with either vehicle (n = 18), SCDi (50 mg/kg, n = 18), PARPi (50 mg/kg, n = 10), or SCDi (50 mg/kg) + PARPi (50mg/kg) (n = 10). *, p < 0.05; **, p < 0.01; ***, p < 0.001.

### Pharmacologic inhibition of SCD induces apoptosis, triggers ER stress, and is associated with DNA damage

Next, we evaluated the functional consequences of pharmacological inhibition of SCD on HBC cells. SCDi decreased the viability of HBC cells in a dose-dependent manner (**Figures 1D, S2A-B**), mainly via apoptosis (**Figures 1E, S2C-D**). Notably, the triple-negative cell lines MDA-MB-231, MDA-BR, HCC1806, and MDA-MB-468 were the most sensitive to SCDi (**Figure S2A**). It is known that the accumulation of SFAs leads to lipotoxicity which activates the endoplasmic reticulum (ER) stress response (24). Consistent with lipotoxicity, we observed a time-dependent increase of ER stress in HBC cells when cells were treated with SCDi (**Figures 1F, S2E-F**). Supplementing cell cultures with 18:1, the major product of SCD, rescued HBC cells from SCDi-induced cell death (**Figure S2G**). These data confirm a critical role for SCD-mediated desaturation in regulating lipotoxicity and influencing cell survival and proliferation.

Recent work from our lab and others has shown a relationship between DNA damage and inhibition of fatty acid synthesis in cancer cells (17,25). To further investigate the mechanism of SCDi-induced cell death in our model, we evaluated DNA damage and repair pathways. DNA double-strand breaks are typically repaired by two major mechanisms: non-homologous end joining (NHEJ) and homology-directed repair (HDR), the latter being mediated by RAD51. Consistent with published work, treatment with SCDi increased the expression of DNA damage marker γ-H2AX and decreased the expression of RAD51 in HBC cells (**Figures 1G and S2H**). Interestingly, when MDA-BR were treated with SCDi and the pan-caspase inhibitor Z-VAD-FMK, which significantly reduced apoptosis (**Figure S2I**), the protein expression of γ-H2AX was decreased (**Figure S2J**), suggesting that DNA damage was caused primarily by engagement of apoptosis. Notably, SCDi-induced downregulation of RAD51 was independent of apoptosis (**Figure S2J**).

Next, we functionally measured DNA damage repair (DDR) using our established bioluminescent repair reporter (BLRR) (23), whereby BLRR cells secrete either Gaussia luciferase (GLuc) when DNA damage is repaired by HDR or Vargula luciferase (VLuc) when DNA damage is repaired through NHEJ (**Figure S2K**). SCDi decreased the ratio HDR/NHEJ in a dose- and time-dependent manner, consistent with a down-regulation of RAD51 (**Figures 1H and S2K**). To explore whether we could use the BLRR reporter to track DDR in vivo, we implanted BLRR-expressing MDA-BR cells into the mammary fat pad, and, upon tumor engraftment, we treated mice with SCDi for 21 days. We measured GLuc and VLuc activity in the blood and confirmed a decreased HDR/NHEJ ratio in mice treated with SCDi (**Figure 1I**). Together, these data suggest that SCDi triggers ER stress, induces DNA damage, and inhibits DDR via RAD51. They also support the idea that SCDi-mediated DNA damage could be an exploitable vulnerability for cancer cells.

### Treatment with SCDi and a PARP inhibitor prolongs survival in a mouse model of BCBM

Inhibitors of Poly(ADP-ribose) polymerase (PARP) exert their anti-tumor effects by inhibiting the repair of DNA single-strand breaks (25). Cancer cells with HDR deficiency are highly sensitive to PARP inhibitors (PARPi) (26–28). Therefore, we hypothesized that impaired HDR following SCD inhibition would sensitize HBC cells to PARPi. We tested this hypothesis using a PARPi-resistant MDA-BR BCBM mouse model (29). MDA-BR were implanted intracranially and, 5 days later, mice were treated with SCDi alone or in combination with the PARPi Niraparib. SCDi treatment led to a significant increase in overall survival, while, expectedly, PARPi alone had no therapeutic effect. However, combining SCDi with PARPi further enhanced the overall survival of BCBM-bearing mice (**Figure 1J**). These data demonstrate the therapeutic efficacy of SCDi in a BCBM mouse model and suggest that SCDi sensitizes BCBM to PARPi by impairing HDR.

### Effect of pharmacological inhibition of SCD in a syngeneic mouse model of BCBM

SCD activity and the balance of SFAs and MUFAs, has been shown to be an important regulator of the inflammatory responses in immune cells (30,31). Therefore, we asked whether pharmacological inhibition of SCD would affect the anti-tumor immune response in a syngeneic mouse model of BCBM. We confirmed that SCDi caused cytotoxicity (**Figure S3A**), which was rescued by oleic acid 18:1 (**Figure S3B**), reduced the FADI (**Figure S3C**), and triggered ER stress (**Figure S3D**) in the mouse mammary cancer cell line EO771. Therefore, as expected, SCDi had the same effects on human and murine mammary cancer cells. Next, immunocompetent mice were intracranially implanted with EO771 cells. Starting at day 4 after implantation, mice received daily doses of vehicle (Control) or SCDi for 14 days (**Figure 2A**). Tumor volume was significantly decreased in mice treated with SCDi (**Figure 2B**). On the day of euthanasia, brain tumors were dissociated, and tumor and immune cells were sorted, and submitted for single-cell RNA sequencing (scRNAseq) (**Figure 2A**). Of note, SCDi-treated tumors showed a higher immune cell/tumor cell ratio when compared to the control (61.2%/29% and 55.6%/33.4%, respectively), reflective of a lower number of tumor cells in SCDi-treated tumors (**Figure S4**).

**Figure 2.**
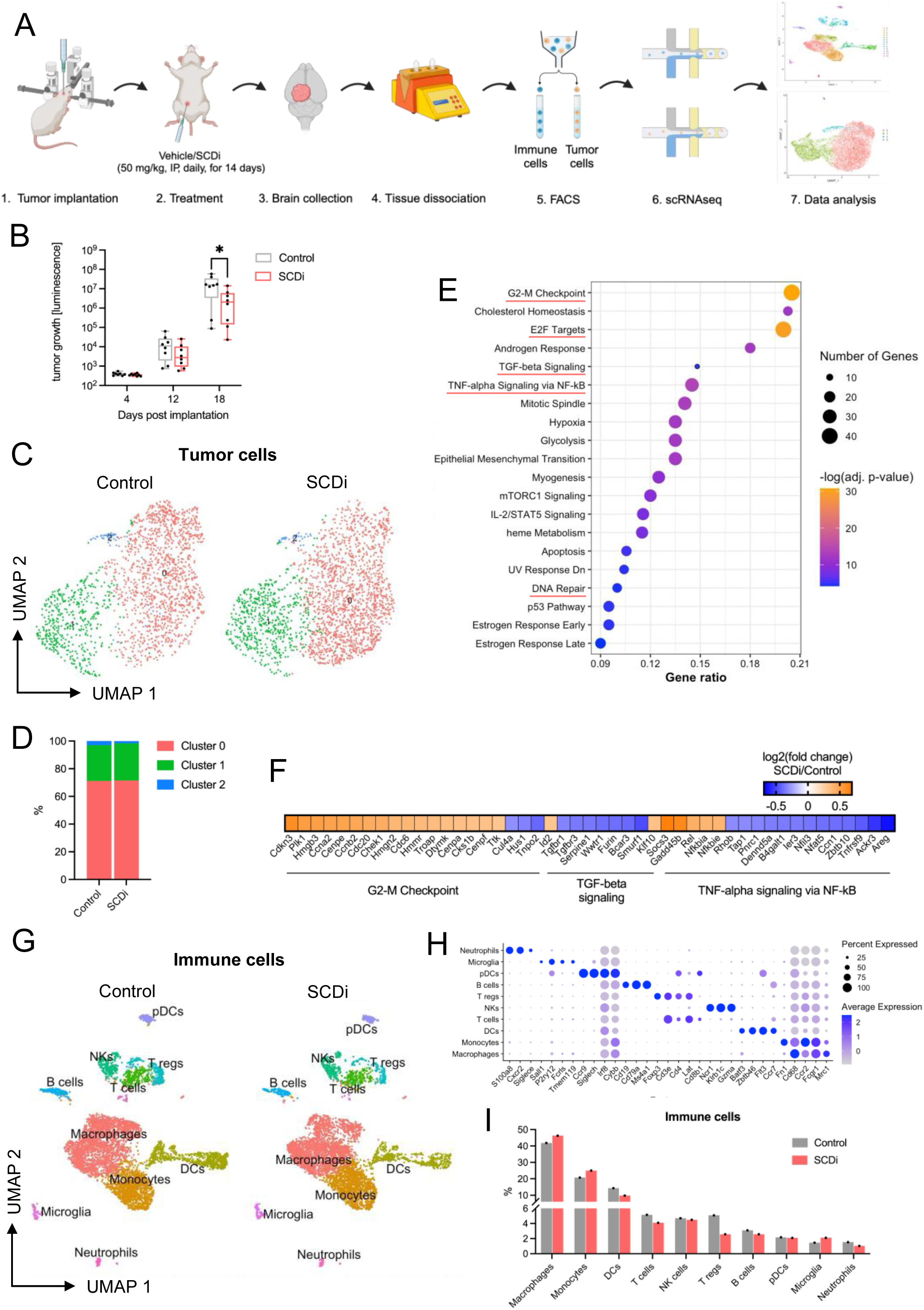
Pharmacological inhibition of SCD remodels the immune cell composition of the tumor microenvironment. (A) Schematic of treatment and tissue processing for single cell RNA sequencing (scRNAseq) in a syngeneic mouse model of breast cancer brain metastasis (BCBM). FACS, fluorescent-activated cell sorting. (B) Luciferase assay from blood collected over time from BCBM-bearing mice to measure tumor volume. (C) UMAP of tumor cells in control and SCDi-treated tumors. (D) Histogram showing proportions of tumor cell subclusters in control and SCDi-treated tumors. (E) Pathway analysis (MSigDB) of differentially expressed genes (DEGs) in tumor cells (SCDi versus control). (F) Heatmap representing DEGs belonging to G2-M checkpoint, TGF-beta signaling, and TNF-alpha signaling via NF-kB pathways in control and SCDi-treated tumors. (G) UMAP of immune cells in control and SCDi-treated tumors. (H) Dot plot of average expression of canonical marker genes for the main immune cell subtypes. (I) Histogram showing cell frequency of immune cells in control and SCDi-treated tumors. *, p < 0.05.

Tumor cells were clustered based on DEGs and visualized on a UMAP plot, that identified 3 main tumor cell subclusters (**Figure 2C**). Due to non-specific gene signatures characterizing these subclusters and to their similar proportions between the treatment groups (**Figure 2D**), we proceeded with the analysis considering all clusters together. We analyzed DEGs in tumor cells between control and SCDi-treated tumors and performed pathway analysis. This analysis revealed a significant dysregulation of the G2/M checkpoint, E2F targets, and DNA repair pathways (**Figure 2E**), with most of the genes belonging to the G2/M checkpoint pathway being upregulated in SCDi-treated tumors (**Figure 2F**). These pathways are activated following DNA damage (32,33). Therefore, dysregulation of these pathways is likely caused by DNA damage upon SCD inhibition, in line with previous data on impaired HDR and elevated DNA damage following treatment with SCDi (**Figures 1G-I and S2H**). The TGF-beta signaling pathway was also significantly altered (**Figure 2E**), with most of the genes (*e.g.*, *Wwtr1*, *Smurf1*, *Serpine1*, *Tgfbr1*) down-regulated in SCDi-treated tumors (**Figure 2F**). This pathway plays a key role in cancer progression and its inhibition is associated with a better outcome (34). Lastly, TNF-alpha signaling via NF-kB was also significantly dysregulated (**Figure 2E**), with most of the genes (*e.g.*, *Nfat5*, *B4galt1*, *Areg*, *Ccn1*, *Ier3*) downregulated in SCDi-treated tumors (**Figure 2F**). Conversely, NF-kB inhibitors *Nfkbia* and *Nfkbie* were upregulated with SCD inhibition, suggesting inhibition of this pathway, as previously shown in ovarian cancer (35).

### Pharmacological inhibition of SCD enhances antigen presentation and interferon signaling of dendritic cells

Unsupervised clustering of CD45^+^ cells based on DEGs identified 10 main cell types, including macrophages, monocytes, dendritic cells (DCs), T cells, natural killer (NK) cells, regulatory T cells (T regs), B cells, plasmacytoid dendritic cells (pDCs), microglia, and neutrophils ( **Figures 2G-I**). Notably, tumors treated with SCDi showed enrichment of monocytes (+21%) and macrophages (+11%), and a decreased percentage of DCs (-31%) and Tregs (-49%), when compared to vehicle-treated tumors (**Figure 2I**).

Next, we analyzed individual immune cell populations, starting with DCs, given their importance in antigen presentation and anti-tumor immunity (36). Cluster analysis of DCs identified two main sub-clusters: mature DCs enriched in immunomodulatory molecules (mregDCs), expressing *Fscn1*, *Mreg*, *Ccr7*, *Stat4*, *Cd200*, *Samsn1*, *Socs2*, and pro-inflammatory antigen-presenting DCs (AP-DCs), expressing *H2-DMb1*, *H2-DMa*, *H2-Ab1*, *Fcer1g*, *Ifitm6*, *Cd209a* (**Figures 3A-B**). MregDCs were recently identified as a novel DC subpopulation in the TME with both immunostimulatory and immunosuppressive functions (37). Compositional analysis revealed a decrease in the percentage of mregDCs and an increase in the percentage of AP-DCs (**Figure 3C**). Based on this, we hypothesized that DCs from SCDi-treated tumors had an enhanced antigen-presenting capacity. We analyzed DEGs in DCs between vehicle- and SCDi-treated tumors and performed a pathway analysis that revealed enrichment of the antigen processing and presentation pathway (**Figure 3D**). In line with our hypothesis, DCs from SCDi-treated tumors showed an upregulation of genes encoding the MHC-II protein subunits (*H2-Aa*, *H2-Ab1, H2-DMa, H2-DMb1, H2-DMb2*, *H2-Eb1*, *H2-Oa*), the invariant chain *Cd74*, a polypeptide that plays a critical role in antigen presentation, facilitating the transport of MHC-II from the ER to the cell surface, and *Ciita*, a master controller of antigen presentation (**Figure 3E**). These data suggests that pharmacological inhibition of SCD is associated with an enhanced antigen presentation capacity of DCs. Moreover, pathway analysis showed a significant dysregulation of pathways related to interferon alpha, interferon gamma, and the inflammatory response (**Figure S5A**). Interestingly, when we analyzed the expression of genes belonging to interferon alpha and gamma response pathways, we found that interferon genes *Isg20*, *Irf7*, and *Ifit3* were upregulated in SCDi-treated tumors compared to the control (**Figure 3E**), suggesting a higher interferon signaling in DCs upon SCD inhibition.

**Figure 3.**
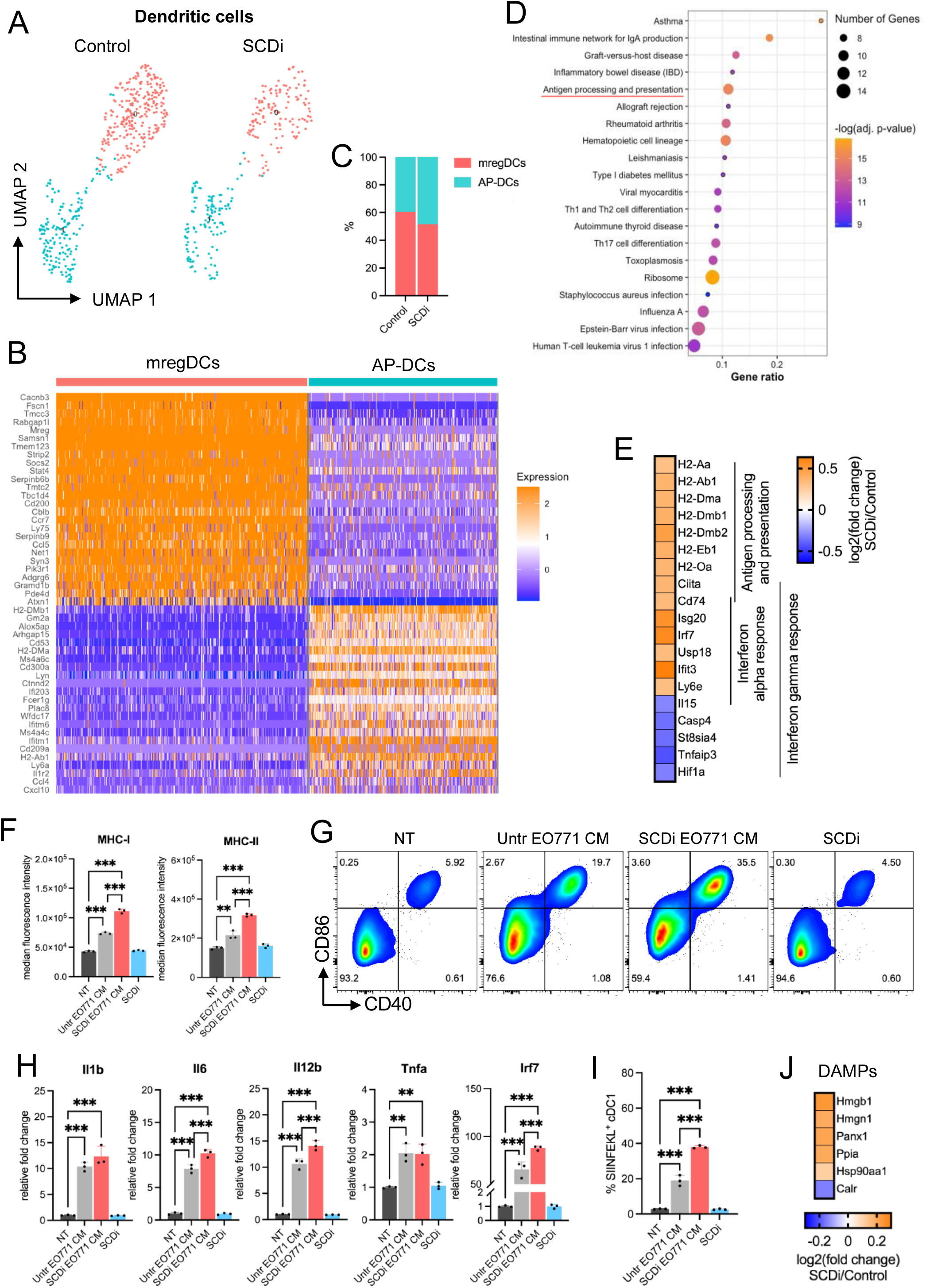
Antigen presentation and interferon signaling of dendritic cells are enhanced in response to pharmacological inhibition of SCD. (A) UMAP of dendritic cells in control and SCDi-treated tumors. (B) Heatmap of the top 20 upregulated genes in the dendritic cell subclusters. mregDCs, mature DCs enriched in immunomodulatory molecules; AP-DCs, antigen-presenting dendritic cells. (C) Histograms representing proportions of dendritic cell subpopulations in control and SCDi-treated tumors (relative to dendritic cells). (D) Pathway analysis (KEGG) of the differentially expressed genes (DEGs) in dendritic cells (SCDi versus control). (E) Heatmap representing DEGs belonging to antigen presentation and processing, interferon alpha, and interferon gamma response pathways in dendritic cells (SCDi versus control). (F) Histograms representing median fluorescence intensity of MHC-I and MHC-II measured by flow cytometry, (G) scatterplot representing the expression of CD40 and CD80 measured by flow cytometry, (H) histograms representing relative fold change of *Il1b*, *Il6*, *Il12b*, *Tnfa*, and *Irf7* measured by qPCR, (I) histograms representing the expression of SIINFEKL+ dendritic cells that were either non-treated (NT), treated with DMSO-treated (Untr) EO771 conditioned medium (CM), with SCDi-treated (SCDi) EO771 CM, or with SCDi alone for 24h. (J) Heatmap representing the gene expression of damage-associated molecular patterns (DAMPs) in tumor cells of SCDi-treated compared to control-treated tumors. **, p < 0.01; ***, p < 0.001.

To validate these results, we generated murine bone marrow-derived dendritic cells (BMDCs) (**Figure S5B-C**). Then, we assessed their activation upon treatment with SCDi- or DMSO-cultured EO771 conditioned medium (CM). Remarkably, the expression of MHC-I, MHC-II (signal I) (**Figure 3F**), CD40, CD80, CD86 (signal 2) (**Figure 3G and S5D**) and *Il1b*, *Il6*, *Il12b*, *Tnfa*, and *Irf7* (signal 3) (**Figure 3H**) were significantly increased in BMDCs exposed to SCDi-treated EO771 CM, when compared to DMSO-treated (Untr) EO771 CM. Next, we employed the ovalbumin model to examine antigen presentation. Briefly, after culture with EO771 CM, DCs were pulsed with the ovalbumin K^b^-binding peptide SIINFEKL for 1h. Staining for OVA_257-264_ peptide (SIINFEKL) loaded onto MHC-I protein showed that stimulation with SCDi-treated EO771 CM increased antigen presentation (**Figure 3I and S5E**). These data may suggest that pharmacological inhibition of SCD induces the release of molecules from cancer cells that function as damage-associated molecular patterns (DAMPs), known to activate immune cells, such as DCs (38). This led us to ask if inhibition of SCD was associated with a higher expression of DAMPs in vivo. Therefore, we used our scRNAseq data and examined the expression of DAMPs in tumor cells. In line with our hypothesis, the expression of several DAMPs (e.g., *Hmgb1, Hmgn1, Panx1, Ppia,* and *Hsp90aa1*) was higher in SCDi-treated tumors when compared to the control (**Figure 3J**). Together, these data suggest that pharmacological inhibition of SCD is associated with an increased MHC and antigen presentation, activation, and interferon signaling of DCs.

### Macrophages from SCDi-treated tumors show higher pro-inflammatory activity

Re-clustering of macrophages identified 3 subpopulations (**Figure 4A**). Analysis of the top 30 highly expressed genes for each subcluster revealed that macrophage subpopulation 0 expressed genes associated with antigen presentation (*H2-Eb1, H2-Ab1, H2-Aa, H2-DMb1, H2-DMa, Cd74)*, and with a pro-inflammatory/anti-tumor M1 phenotype (*Il-1β* and *Cxcl9*). Macrophage subpopulation 1 highly expressed *Folr2*, a gene that has been associated with CD8^+^ T cell infiltration and favorable prognosis in HBC (39), and many cytokines (*Ccl2, Ccl4, Ccl6, Ccl7, Ccl8, Ccl9, Cxcl10*). Macrophage subpopulation 2 expressed genes not indicative of a specific status (**Figure 4B**). Compositional analysis revealed an increase of *Folr2^+^*/cytokines-expressing macrophages in SCDi-treated tumors (**Figure 4C**). Next, we analyzed the DEGs between vehicle- and SCDi-treated macrophages. We performed a pathway analysis, which revealed an enrichment of pathways associated with interferon alpha, interferon gamma, and inflammatory response (**Figure 4D**). The majority of DEGs belonging to these pathways (*e.g.*, *Ifitm3, Isg20, Psme1, Irf7, Isg15, Gbp2, Ifit3, Ccl7, Fgl2*) were upregulated in macrophages from SCDi-treated tumors (**Figure 4E**), showing higher interferon signaling in these tumors. These data suggest an enhanced pro-inflammatory/anti-tumor activity of macrophages upon pharmacological inhibition of SCD.

**Figure 4.**
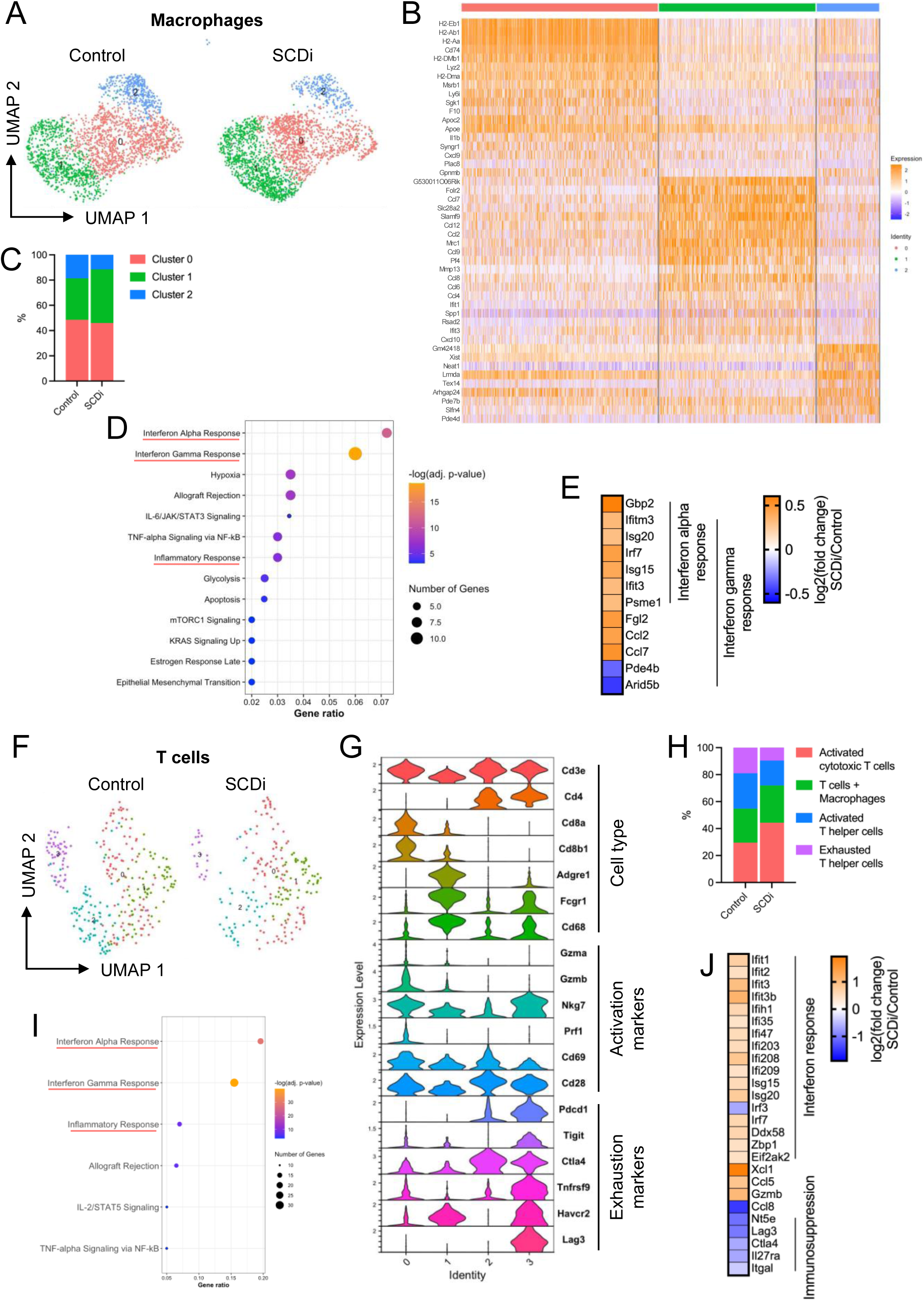
Inhibition of SCD enhances interferon signaling in macrophages and T cells and decreases T cell exhaustion in the tumor microenvironment. (A) UMAP of macrophages in control and SCDi-treated tumors. (B) Heatmap of the top 20 upregulated genes in the macrophage subclusters. (C) Histograms representing proportions of macrophage subpopulations in control and SCDi-treated tumors. (D) Pathway analysis (MSigDB) of the differentially expressed genes (DEGs) in macrophages (SCDi versus control). (E) Heatmap representing the DEGs belonging to the interferon alpha and gamma response pathways in macrophages (SCDi versus control). (F) UMAP of T cells in control and SCDi-treated tumors. (G) Violin plots showing the expression of cell type, activation, and exhaustion markers in T cells subclusters. Cluster 0 represents activated cytotoxic T cells, cluster 1 represents T cells + Macrophages, cluster 2 represents activated T helper cells, and cluster 3 represents exhausted T helper cells. (H) Histograms representing proportions of T cells subclusters in control and SCDi-treated tumors. (I) Pathway analysis (MSigDB) of the DEGs in T cells (SCDi versus control). (J) Heatmap representing a subset of relevant DEGs in T cells (SCDi versus control).

### Pharmacological inhibition of SCD triggers anti-tumor T cell responses in the TME

We also analyzed the T cell gene expression programs in response to SCDi. Unsupervised clustering of T cells identified 4 main subpopulations: activated cytotoxic T cells (*CD3e, CD8a, Cd8b1, Gzmb, Nkg7, Prf1, Cd69, Cd28*), a subpopulation composed of T cells (*Cd3e*) and contaminating macrophages (*Adgre1, Fcgr1, Cd68*) that we named T cells + macrophages, activated T helper cells (*Cd3e, Cd4, Cd69, Cd28*), and exhausted T helper cells (*Cd3e, Cd4, Pdcd1, Tigit, Ctla4, Tnfrsf9, Havcr2, Lag3*) (**Figures 4F-G**). Compositional analysis revealed a higher proportion of activated cytotoxic T cells and a lower proportion of exhausted T helper cells in SCDi-treated tumors compared to the control (**Figure 4H**). Pathway analysis of the DEGs between control- and SCDi-treated tumors showed significant enrichment of interferon alpha, interferon gamma, and inflammatory response pathways (**Figure 4I**). Analysis of specific gene signatures showed an upregulation of many interferon genes (*e.g.*, *Ifit1, Ifit2, Ifit3, Isg20, Irf7*), chemokines (*Xcl1* and *Ccl5*) known to play an important role in T cell activation and cDC recruitment in the TME (40), and *Gzmb*, a potent cytotoxic protease used by cytotoxic T cells to kill cancer cells, in SCDi-treated tumors compared to the control (**Figure 4J**). We also noted downregulation of *Ccl8* in SCDi-treated tumors. Ccl8 is associated with Tregs recruitment (41), providing additional evidence for decreased recruitment and presence of intratumoral Tregs upon SCD inhibition, as shown in compositional analysis of different immune cell populations (**Figure 2I**). Pharmacological inhibition of SCD also decreased the expression of *Lag3* and *Ctla4*, which are associated with T cell exhaustion (**Figure 4J**). These data suggest that SCDi induces a higher interferon response in T cells and decreases the immunosuppressive effects of Tregs and exhausted T cells to support anti-tumor T cell responses.

### CellChat analysis shows stronger interactions via MHC-I and weaker immunosuppressive PD-1:PD-L1/PD-L2 and PVR/TIGIT axes interactions upon SCD inhibition

The transcriptomic changes in immune cells in response to pharmacological inhibition of SCD led us to perform cell-to-cell interaction analysis across all immune cell subpopulations using CellChat. Initially, we examined the signaling pathways of all receptor-ligand interactions between the immune cell populations (**Figure 5A**). SCDi-treated tumors showed stronger interactions (thicker lines) between all APCs (AP-DCs, mregDCs, monocytes, macrophages, neutrophils, microglia, pDCs, and B cells) and T/NK cells via MHC-I (**Figures 5B and S6A**), consistent with higher antigen presentation through MHC-I when SCD is inhibited, in line with our functional DC studies (**Figure 3F**). Additionally, SCDi-treated tumors showed similar interactions between APCs (AP-DCs, macrophages, and B cells) and T cells via MHC-II (**Figures 5C and S6B**). Moreover, in line with our pathway analyses, CellChat showed weaker interactions via PD-L1/PD-L2 immune checkpoints (**Figures 5D-E and S6C-D**) and via PVR/TIGIT (**Figures 5F-G and S6E-F**). The latter is an immunosuppressive axis that leads to an anti-inflammatory, non-proliferative, non-cytotoxic profile of T cells (42,43). Also, the phosphoprotein 1 (SPP1) signaling pathway, previously associated with immunosuppression and poor prognosis in lung cancer,(44) showed decreased interactions, particularly between macrophages and other immune cells (AP-DCs, T cells, NKs, and B cells) in SCDi-treated tumors (**Figure S6G**). Treatment with SCDi also resulted in stronger interactions via the LCK signaling pathway (**Figure S6H**), which is important for T cell activation.(45) We also found stronger interactions via the Notch pathway, Nkg2d, and Il12 upon SCD inhibition (**Figure 5A**). Together, these data support a higher activation status via MHC-I and LCK pathways and a diminished level of immunosuppression via PD-L1/PD-L2, and PVR/TIGIT in response to SCDi, likely contributing to an enhanced anti-tumor T cell response.

**Figure 5.**
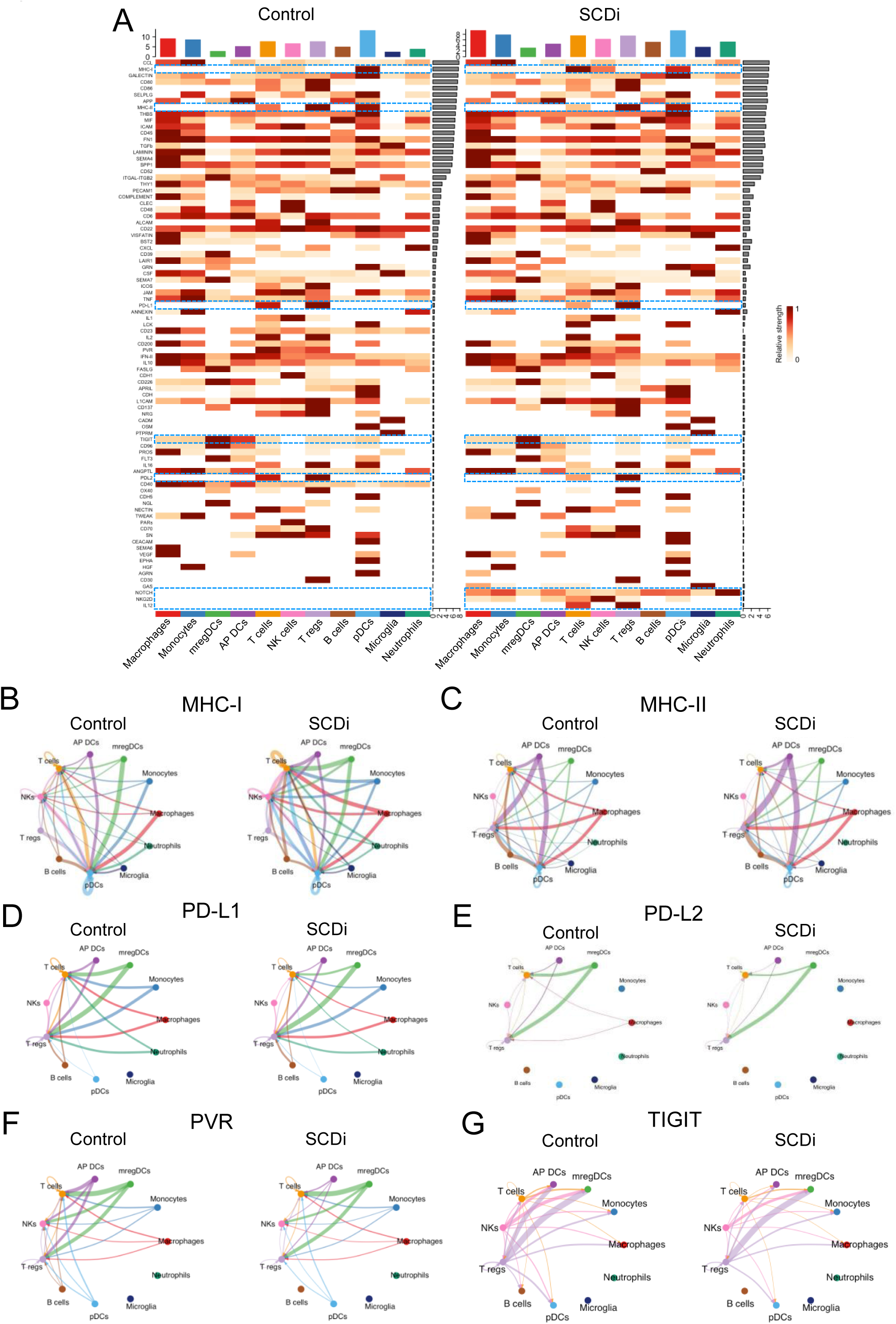
Interactions via PD-1:PD-L1/PD-L2 and PVR/TIGIT immunosuppressive axes are weakened in response to inhibition of SCD. (A) Overall signaling patterns between CellChat-curated pathways and cell clusters. The relative strength reflects the sum of all normalized interactions in each pathway. Inferred (D) MHC-I, (E) MHC-II, (F) PD-L1, (G) PD-L2, (H) PVR, (I) TIGIT signaling networks among the cell populations represented by the nodes. The thickness of lines connecting two cell populations represent the interaction strength (*i.e.*, thicker lines represent stronger interactions).

## Discussion

This study explored the therapeutic efficacy of SCDi on BCBM and the role of pharmacological inhibition of SCD on the TME. First, we tested the effect of SCDi on different HBC cell lines. Jin and co-authors showed high brain tropism of the triple-negative breast cancer cell lines MDA-MB-231 and HCC1806 in mouse xenografts (13). In our study, these two cell lines, and the brain-tropic cell line MDA-BR, showed the highest sensitivity to SCDi, suggesting a high efficacy of this inhibitor on brain metastatic cells. PARP inhibitors, currently being tested for the treatment of BCBM (46), showed increased potency in BRCA-deficient compared to BRCA-wild-type tumors (47). Additionally, high RAD51 expression predicted poor response to PARPi (48). As a result, it was proposed that the combination of RAD51 and PARP inhibitors was effective in BRCA-wild-type breast cancer cells (29). In line with a downregulation of RAD51 and impaired HDR following SCD inhibition in vitro and in vivo, we found that combining SCDi with PARPi exhibited greater anti-tumor activity in a BRCA-wild type mouse model of BCBM. These results suggest that tumor cells could become vulnerable to PARPi upon SCD inhibition and support further testing of SCD and PARP inhibitors as a combination therapy for BCBM and other PARPi-resistant tumors.

Fatty acid metabolism reprogramming has been shown to affect the proliferation, differentiation, and activation of immune cells in the TME (49). Consequently, a growing body of research has explored the role of fatty acid metabolism in immune cells to develop new anti-tumor therapeutic strategies. When we analyzed the effect of SCD inhibition on the immune cells, we found prominent changes with clinical relevance if translated to humans. We found evidence for increased DC activation upon SCD inhibition. Multiple mechanisms might explain this. For example, following DNA damage, tumor cells accumulate DNA in the cytosol that functions as an immunogenic danger signal, ultimately activating the expression of type I interferon-stimulated genes (ISGs) (50). Moreover, dying tumor cells shed DNA that activates DCs via toll-like receptor 3 (TLR3), TLR9, and cGAS/STING, as previously reported (50,51). Lastly, other molecules, known as DAMPs, released by dying cancer cells activate other TLRs (*i.e.*, TLR2 and TLR4), ultimately leading to a higher immune cell activation (38). We also found higher interferon signaling in the TME of SCDi-treated tumors. Interferons are key components of immune cell activation.(52) They act on tumor cells to exert a direct anti-tumor effect, inducing apoptosis (53). They can also regulate the activity of many immune cell types, leading to a balanced anti-tumor immune response. For example, they can activate DCs to enhance antigen presentation to T cells and recruit CD4^+^ Th1 and CD8^+^ T cells to the TME (54). Also, interferons enhance macrophage activity by polarizing monocytes into mature pro-inflammatory anti-tumor M1 macrophages (53,55). Moreover, interferons regulate the cytotoxic and anti-tumor activity of NK and T cells to exert a more effective anti-tumor immune response (56).

In line with heightened inflammatory responses in SCDi-treated tumor-bearing mice, we observed a shift in the balance of effector and immunosuppressive T cells in the TME. Previous studies reported that SCD inhibition was associated with an enhanced proliferation of CD8^+^ T cells in various mouse tumor models (22), and with a greater T effector function in an LCMV infection model (19). We found that SCDi-treated tumors were associated with reduced T cell exhaustion, diminished recruitment of Tregs, and dampened signaling through immunosuppressive pathways, such as the PD-L1/PD-L2 and the PVR/TIGIT axes. T cell exhaustion is a hypofunctional T cell state characterized by a decreased anti-tumor effect (57). Tregs have an immunosuppressive role in inflammation and can suppress anti-tumor immunity (58). The PD-L1/PD-L2 signaling pathway can inhibit the activation and effector function of T cells, ultimately driving tumor immune escape (59). Similarly, the immunosuppressive PVR/TIGIT pathway is currently being investigated as a target for immunotherapy (43). We think this effect is likely a combination of intrinsic effects that SCDi has on T cell function and the impact of SCDi on antigen-presenting cells (*i.e.*, DCs and macrophages) in the TME. We also propose that the impact of SCDi on DNA damage and tumor cell viability further contributes. Overall, these data suggest that SCD inhibition is a potential strategy for turning “cold” immune-excluded tumors into “hot” immune-inflamed ones. Brain tumors are considered “cold” tumors, characterized by an immunosuppressive TME, and a poor response to immunotherapy, partly due to the highly complex brain microenvironment (60,61). The premise of developing a strategy that prompts an immune-inflamed TME and potentially bolsters immunotherapy is promising. As such, we speculate that combining SCDi with immune checkpoint inhibitors could be a strategy to treat BCBM that warrants further investigation.

There are limitations to this study. First, the efficacy of SCDi was explored using one xenograft (MDA-BR) and one syngeneic (EO771) mouse model. Further studies with additional mammary cancer cell lines are needed to corroborate our results. Also, the molecular mechanisms by which DCs show higher antigen-presenting ability and T cells show dampened exhaustion upon pharmacological inhibition of SCD remain to be elucidated. Moreover, an in-depth analysis of the role of SCD in distinct immune cell populations and an in vitro validation of the enhanced anti-tumor response of T cells upon SCD inhibition are limiting. Yet, whether the changes in the TME are due to a direct effect of SCD inhibition on immune cells or solely to the indirect effect of cancer cell death remains to be elucidated. Lastly, whether SCDi can be used prophylactically in patients who undergo surgical resection of the primary tumor to prevent the development of brain metastases still needs to be explored and might be an interesting approach for future studies.

In conclusion, this study highlights the dual role of pharmacological inhibition of SCD in BCBM mouse models. We show that a brain-penetrant SCDi triggers lipotoxicity and impaired DNA damage repair in tumor cells while promoting an anti-tumor response in the TME. Further studies are needed to explore whether combining inhibition of SCD with immunotherapy might be a novel therapeutic strategy to treat these incurable tumors.

## Materials and methods

### Cell lines

Human breast cancer cell lines MDA-MB-231, MCF7, HCC1806, MDA-MB-468, and BT474, and mouse mammary cancer cell line EO771 were purchased from American Type Culture Collection (ATCC; Manassas, VA, USA). Brain tropic MDA-MB-231-BR (MDA-BR) cells were kindly provided by Dr Patricia S. Steeg (National Cancer Institute, Bethesda, MD). All cells were cultured at 37°C in a 5% CO_2_ humidified incubator. The culture medium for MDA-MB-231, MCF7, MDA-MB-468, and MDA-BR cells was Dulbecco’s Modified Essential Medium (DMEM, Corning) supplemented with penicillin (100 units/mL), streptomycin (100 mg/mL) (P/S) (Corning) and 10% fetal bovine serum (FBS; Atlanta Biologics). The culture medium for HCC1806 and BT474 cells was Roswell Park Memorial Institute (RPMI, ATCC) supplemented with P/S (Corning) and 10% FBS (Atlanta Biologics). The culture medium for EO771 cells was DMEM supplemented with P/S (Corning), 10% FBS (Atlanta Biologics), and 20 mM HEPES (Thermo Fisher Scientific).

### Reagents/test inhibitors

For *in vitro* studies, YTX-7739 (SCDi), Niraparib (MK-4827, Selleckchem), and Z-VAD-FMK (Selleckchem) were dissolved in DMSO. For *in vivo* studies, YTX-7739 was dissolved by sonication (2-3 minutes), heating (3 minutes at 60°C), and vortexing in 0.5% methylcellulose + 0.2% Tween80 in sterile saline solution. Niraparib was prepared as per manufacturer’s instructions. Bovine Serum Albumin (BSA)-conjugated oleate (18:1) monounsaturated fatty acid complex was obtained from Cayman chemical.

### 13C-Glucose labeling experiments

1-4 x 10^5^ cells were plated in 12-well plates in 10% FBS media. When attached, cells were treated with DMSO/SCDi in serum-free glucose-free media, supplemented with 2.25g/L of unlabeled D-glucose (Millipore Sigma) and 2.25g/L of ^13^C_6_-glucose (Cambridge Isotope Laboratories). Cells were collected and counted using the Cellometer K2 Cell Counter (Nexcelom). Then, 1-3 x 10^5^ cells were resuspended in 3M methanolic guanidine HCl and immediately transferred to glass tubes for derivatization. Samples were prepared alongside internal standard curve samples made up of FAMEs (Nu-Chek Prep). Isotopomer spectral analysis (ISA) was conducted using an Agilent 5975C mass spectrometer coupled to a 7890 Gas Chromatograph. Data were normalized by total cell count.

### Shotgun lipidomics

4×10^5^ MDA-BR cells per well of a 6-well plate were plated and treated with DMSO/SCDi (1 μM) for 48 hours in 4 replicates. Cells were then collected and counted using the Cellometer K2 Cell Counter (Nexcelom). Lipids were extracted from 1×10^6^ cells as previously described (17). Shotgun lipidomic analysis was performed as previously described (17). Quantitative values of lipid species were normalized to cell counts. Heatmaps and Principal Component Analysis (PCA) plots were generated using Clustvis (https://biit.cs.ut.ee/clustvis/). Nipals PCA was used to calculate principal components (PCs) on PCA.

### Lentiviral vectors and transduction

CSCW-GLuc-IRES-GFP lentiviral vector expressing Gaussia Luciferase (GLuc) and green fluorescent protein (GFP) was used. To transduce cancer cell lines, cells (2-4 x 10^5^) were seeded in a 6-well plate. When cells were attached, polybrene (10 μg/ml; Millipore Sigma) and 10-30 μl of lentivirus were added to each well. Cells were then spun at 1,800 rpm at 4°C for 1.5 hours and incubated at 37°C. After 12-14 hours, the medium was replaced with fresh medium. GFP-positive cells were then selected by Fluorescent Activated Cell Sorting (FACS).

### In vitro assays

Cell viability assays were performed using CellTiter-Glo 2.0 (Promega). Apoptosis was measured using Caspase-Glo 3/7 (Promega). Early apoptosis and late apoptosis/necrosis were measured using Alexa Fluor 488 Annexin V/Dead Cell Apoptosis Kit (Thermo Fisher Scientific). These reagents were used as indicated by the manufacturer.

### Protein extraction and western blotting

Radioimmunoprecipitation assay buffer (RIPA) buffer (Thermo Fisher Scientific) supplemented with protease and phosphatase inhibitors (Thermo Fisher Scientific) was used to lyse cells and extract proteins, as recommended by the manufacturer. Protein concentration was calculated using the Pierce BCA Protein Assay Kit (Thermo Fisher Scientific), following manufacturer’s instructions, and 10-30 μg of proteins were loaded and resolved on either a 10% or 4-12% Bolt Bis-Tris gel (Thermo Fisher Scientific), transferred to a nitrocellulose membrane (Bio-Rad), and then incubated with the indicated antibodies. Proteins were then detected with SuperSignal West Pico Chemiluminescent Substrate (Thermo Fisher Scientific). The following antibodies were used: PARP (CST #9532), phospho-eIF2α (CST #3597), ATF-4 (CST #11815), Phospho-Histone H2A.X (γ-H2AX) (CST #9718), anti-rabbit IgG, HRP-linked (CST #7074) and Anti-mouse IgG, HRP-linked (CST #7076), GAPDH (Santa Cruz #sc-47724), beta actin (Abcam #ab227387), Rad51 (BIOSS Antibodies #BSM-51402M).

### RNA extraction and quantitative PCR

RNA was extracted using the Quick-RNA MiniPrep kit (Zymo Research). RT-PCR was performed using the High-Capacity cDNA Reverse Transcription Kit (Thermo Fisher Scientific). qPCR was performed using the PowerUP SYBR Green Master Mix (Thermo Fisher Scientific) using QuantStudio 3 System. All reagents were used as recommended by the manufacturers. Relative mRNA expression was calculated using the 2^-ΔΔCT^ method. Beta-actin or HPRT1 were used as housekeeping genes. Primer sequences are reported in Table S1.

### In vivo mouse models

Animal experiments were approved by the Massachusetts General Hospital Subcommittee on Research Animal Care or by the University of California, Los Angeles Animal Research Committee. For orthotopic mouse models, female athymic nude mice were injected into the mammary fat pad with 1×10^6^ MDA-BR cells in a 100μl mixture composed of Matrigel (50μl; BD Matrigel 10mg/ml, BD Biosciences) and PBS (50μl). For intracranial mouse models, female mice were intracranially injected with 5×10^4^ MDA-BR cells (athymic nude mice) or 5×10^2^ EO771 expressing Luc-GFP (C57BL/6J) in PBS (2μL) as previously described (17). For *in vivo* studies, YTX-7739 (100μl) was administered by intraperitoneal injections, daily. Niraparib (100μl) was administered by oral gavage, daily. Control groups were administered with 100μl of vehicle (0.5% methylcellulose + 0.2% Tween80 in saline solution).

### Bioluminescence BLRR assay to monitor DNA damage repair

The bioluminescent repair reporter (BLRR) system was used to track DNA damage repair as previously described (23). To detect BLRR activity *in vitro*, cells were plated in a 96-well plate and treated with DMSO/SCDi. All luminescence results were normalized to cell viability measured using CellTiter-Glo2.0. To detect BLRR activity *in vivo*, 1×10^6^ MDA-BR cells were injected into the mammary fat pad of female athymic nude mice. Mice were treated with vehicle/SCDi, as described. On the day of euthanasia, blood was collected into a K_2_-EDTA blood collection tube and spun at 2,000rcf for 15 minutes. 25μl of plasma were transferred into a white 96-well plate to measure luminescence, as described above.

### Tissue digestion

Neural Tissue Dissociation Kit (P) (Miltenyi Biotec) was used to digest the brains into a single cell suspension, according to manufacturer’s instructions. Myelin was removed using anti-myelin beads (Miltenyi Biotec), following manufacturer’s instructions. The final single cell suspension was resuspended in 1X PBS w/o calcium and magnesium, supplemented with 2nM EDTA (Thermo Fisher Scientific) and 0.5% Bovine Serum Albumin (BSA; Millipore Sigma), before cell staining and FACS.

### scRNAseq

For scRNAseq, 3 mice per experimental group were pooled into 1 sample prior to FACS-sorting to account for inter-animal variability. Each brain pool was sorted into 2 cell populations: tumor cells (DAPI^-^/CD45^-^/GFP^+^) and immune cells (DAPI^-^/CD45^+^/GFP^-^). Within 30 minutes after sorting, 3’ scRNA-seq libraries were generated using a 10X Genomics Chromium controller. Libraries were then sequenced on a NovaSeq 6000 platform using a NovaSeq S2 flow cell (Illumina). Data analysis was performed in R studio (v1.3.1073) using Seurat (v4.1.1). Cells with less than 1000 detected genes, UMI numbers <300 or with >2% mitochondrial-derived UMI counts were excluded from the analysis. Raw Fastq data were processed using CellRanger v3.0.2. Cell clusters were identified with the FindClusters function and visualized on uniform manifold approximation and projection (UMAP) plots using the top 30 dimensions for the analysis. Cell types were annotated based on corresponding literature. Differentially expressed genes (DEGs) of cell subgroups were recognized by the FindAllMarker function, using |logFC| = 0.3 and adjusted p-value <0.05 as the cut-off criteria. Pathway on DEGs was performed using enrichr, KEGG_2019_Mouse and HDSigDB_Mouse_2021 packages.

### Bone marrow-derived dendritic cell culture

To generate bone marrow-derived dendritic cells (BMDCs), bone marrow progenitors were resuspended at 1.5×10^7^ cells in 10mL of BMDC medium (RPMI 1640, 1% penicillin-streptomycin, 10% FBS, 1X Glutamax, 200mM sodium pyruvate, 25mM HEPES, 1% MEM-NEAA, 0.5% sodium bicarbonate, 0.01% 55mM 2-mercaptoethanol supplemented with 200ng/mL FLT3-L and 5ng/mL GM-CSF), and seeded in 10cm non-TC treated culture dishes and incubated at 37°C, 5% CO_2_. On day 5, 5mL of fresh media were added to the cells. On day 9, media was replaced, and cells were used for experiments on day 15.

### Collection of EO771 conditioned medium and treatment of BMDCs

To generate EO771 conditioned medium (CM), EO771 cells were seeded at a density of 1.5×10^5^ cells per well of a 6-well plate in complete medium. The next day, the medium was replaced with DMEM/F12 + 1% penicillin-streptomycin supplemented with either 0.1% DMSO (Untr EO771 CM) or 1μM SCDi (SCDi EO771 CM). After 48h, the medium was centrifuged at 300rcf for 5 minutes to remove cell debris. For CM treatment, 5×10^5^ BMDCs were seeded in a non-TC treated 24-well plate in 250μL complete medium + 250μL EO771 CM. BMDCs were analyzed after 48h incubation. For the antigen presentation assay, BMDCs were exposed to EO771 CM for 24h, and then pulsed with 1μM OVA_257-264_ (Invivogen) for 1h. Cells were washed three times with PBS and stained for flow cytometry analysis.

### Flow Cytometry

Non-specific binding was blocked by incubating cells with TruStain FcX (anti-mouse Cd16/32, BioLegend) for 5-10 minutes. Cells were then stained with the antibodies for 30 minutes on ice in the dark. Cells were then washed with PBS + 2%FBS + 2mM EDTA (FACS buffer), centrifuged at 300rcf for 5 minutes, and resuspended in FACS buffer. DAPI was added to the single cell suspension at a final concentration of 1 μg/ml to stain dead cells. Cells were then sorted using a BD FACSAria II cell sorter. Cells for single cell RNA sequencing were sorted into 100% FBS. The following antibodies were purchased from BioLegend: BV785-anti-CD45, BV711-anti-CD11c, ACP/Cy7-anti-I-A/I-E, BV650-anti-XCR1, AF647-anti-H-2Kb/H-2Db, PE/Cy7-anti-CD40, PE/Dazzle594-anti-CD80, BV421-anti-CD86, APC-anti-H-2Kb bound to SIINFEKL.

### Statistical analysis

Statistical analysis was performed using Prism 10. Statistical tests are reported in the corresponding figure legends. Level of significance was fixed as p<0.05.

### Data availability

The data presented in this manuscript will be made available upon reasonable request to the corresponding authors.

## Acknowledgments

We are grateful to Yumanity Therapeutics for kindly providing YTX-7739. We would like to thank the Flow Cytometry Core at UCLA and the Applied Genomics, Computational & Translational Core at Cedars-Sinai Medical Center. We would like to thank Christopher Tse, David Nathanson, and Semer Maksoud for their help and Xandra Breakefield for her support. Graphics were created with Biorender (Biorender.com).

## Author contributions

Conceptualization: A.S., S.J.B., and C.E.B.; Methodology: A.S., S.J.B., and C.E.B.; Validation: A.S., G.G., K.M.E., K.J.W., S.J.B., and C.E.B.; Formal Analysis: A.S., G.G., and B.S.; Investigation: A.S., K.M.E., K.K., Y.Q., J.E.H., K.J.W., and V.Z.; Resources: K.J.W., S.J.B., and C.E.B.; Data Curation: A.S. and G.G.; Writing Original Draft: A.S., S.J.B., and C.E.B.; Writing – Review & Editing: G.G., K.M.E., K.K., Y.Q., J.E.H, K.J.W., B.S., and V.Z.; Visualization: A.S. and G.G.; Supervision: S.J.B. and C.E.B.; Project Administration: S.J.B. and C.E.B.; Funding Acquisition: S.J.B., and C.E.B.

