## Supplementary material for "Targeting SCD triggers lipotoxicity of cancer cells and enhances anti-tumor immunity in breast cancer brain metastasis mouse models": Figure S

Sammarco et al.

This PDF file includes:

Materials and Methods
Figures S1 to S6
Table S1

### Figure S1

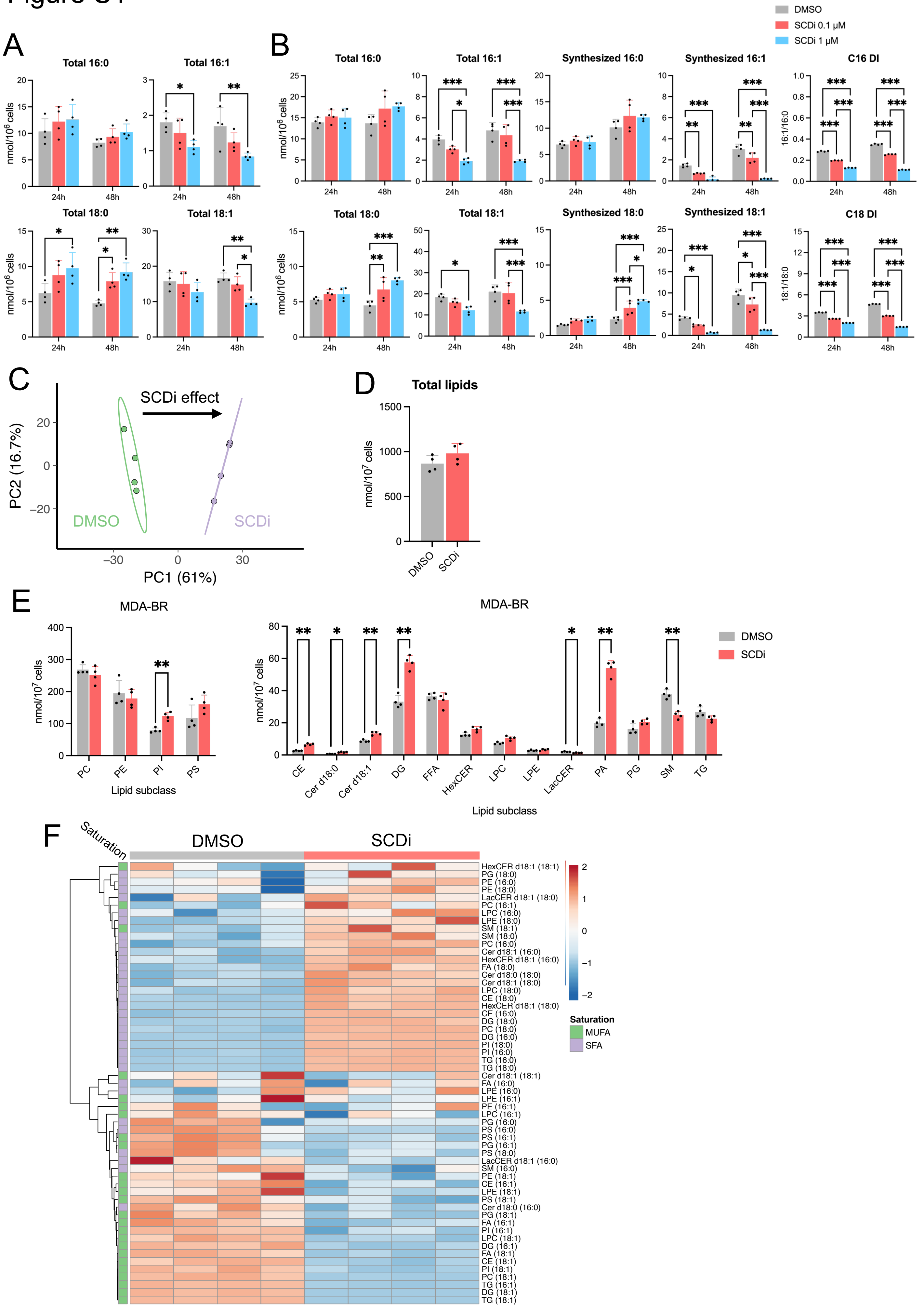

DMSO
SCDi 0.1 µM
SCDi 1 µM
A
Total 16:0
Total 16:1
Total 18:0
Total 18:1
nmol/10^6 cells
24h
48h
B
Total 16:0
Total 16:1
Synthesized 16:0
Synthesized 16:1
C16 DI
16:1/16:0
Total 18:0
Total 18:1
Synthesized 18:0
Synthesized 18:1
C18 DI
18:1/18:0
C
SCDi effect
PC2 (16.7%)
PC1 (61%)
DMSO
SCDi
D
Total lipids
nmol/10^7 cells
DMSO
SCDi
E
MDA-BR
nmol/10^7 cells
PC
PE
PI
PS
Lipid subclass
CE
Cer d18:0
Cer d18:1
DG
FFA
HexCER
LPC
LPE
LacCER
PA
PG
SM
TG
DMSO
SCDi
F
Saturation
DMSO
SCDi
HexCER d18:1 (18:1)
PG (18:0)
PE (16:0)
PE (18:0)
LacCER d18:1 (18:0)
PC (16:1)
LPC (16:0)
LPE (18:0)
SM (18:1)
SM (18:0)
PC (16:0)
Cer d18:1 (16:0)
HexCER d18:1 (16:0)
FA (18:0)
Cer d18:0 (18:0)
Cer d18:1 (18:0)
LPC (18:0)
CE (18:0)
HexCER d18:1 (18:0)
CE (16:0)
DG (18:0)
PC (18:0)
DG (16:0)
PI (18:0)
PI (16:0)
TG (16:0)
TG (18:0)
Cer d18:1 (18:1)
FA (16:0)
LPE (16:0)
LPE (16:1)
PE (16:1)
LPC (16:1)
PG (16:0)
PS (16:0)
PS (16:1)
PG (16:1)
PS (18:0)
LacCER d18:1 (16:0)
SM (16:0)
PE (18:1)
CE (16:1)
LPE (18:1)
PS (18:1)
Cer d18:0 (16:0)
PG (18:1)
FA (16:1)
PI (16:1)
LPC (18:1)
DG (16:1)
FA (18:1)
CE (18:1)
PI (18:1)
PC (18:1)
TG (16:1)
DG (18:1)
TG (18:1)
Saturation
MUFA
SFA

**Figure S1. SCDi reshapes the lipid composition of breast cancer cells.**

(A-B) Gas chromatography/mass spectrometry of (A) HCC1806 and (B) MDA-MB-468 treated with SCDi for 24h and 48h, measuring total and synthesized palmitic acid (16:0), palmitoleic acid (16:1), stearic acid (18:0), and oleic acid (18:1). On the right, the desaturation index (DI - ratio 16:1/16:0 and 18:1/18:0) is shown. Mean □ SD (Two-way ANOVA, Tukey's test).

(C) Principal component analysis of individual lipids quantified using shotgun lipidomics from MDA-BR cells treated with DMSO or SCDi (1 □M) for 48h. PC1, principal component 1; PC2, principal component 2.

(D) Histograms representing the amount of total lipids quantified using shotgun lipidomics from MDA-BR cells treated with DMSO or SCDi (1 □M) for 48h.

(E) Histograms representing the amounts of 17 lipid subclasses quantified using shotgun lipidomics from MDA-BR cells treated with DMSO or SCDi (1 □M) for 48h. Mean □ SD (Student's t test).

(F) Heat map representing the compositional analysis of individual lipids containing palmitic acid (16:0), palmitoleic acid (16:1), stearic acid (18:0), and oleic acid (18:1) quantified using shotgun lipidomics of MDA-BR treated with SCDi (1□M) for 48h. *, $p < 0.05$; **, $p < 0.01$.

### Figure S2

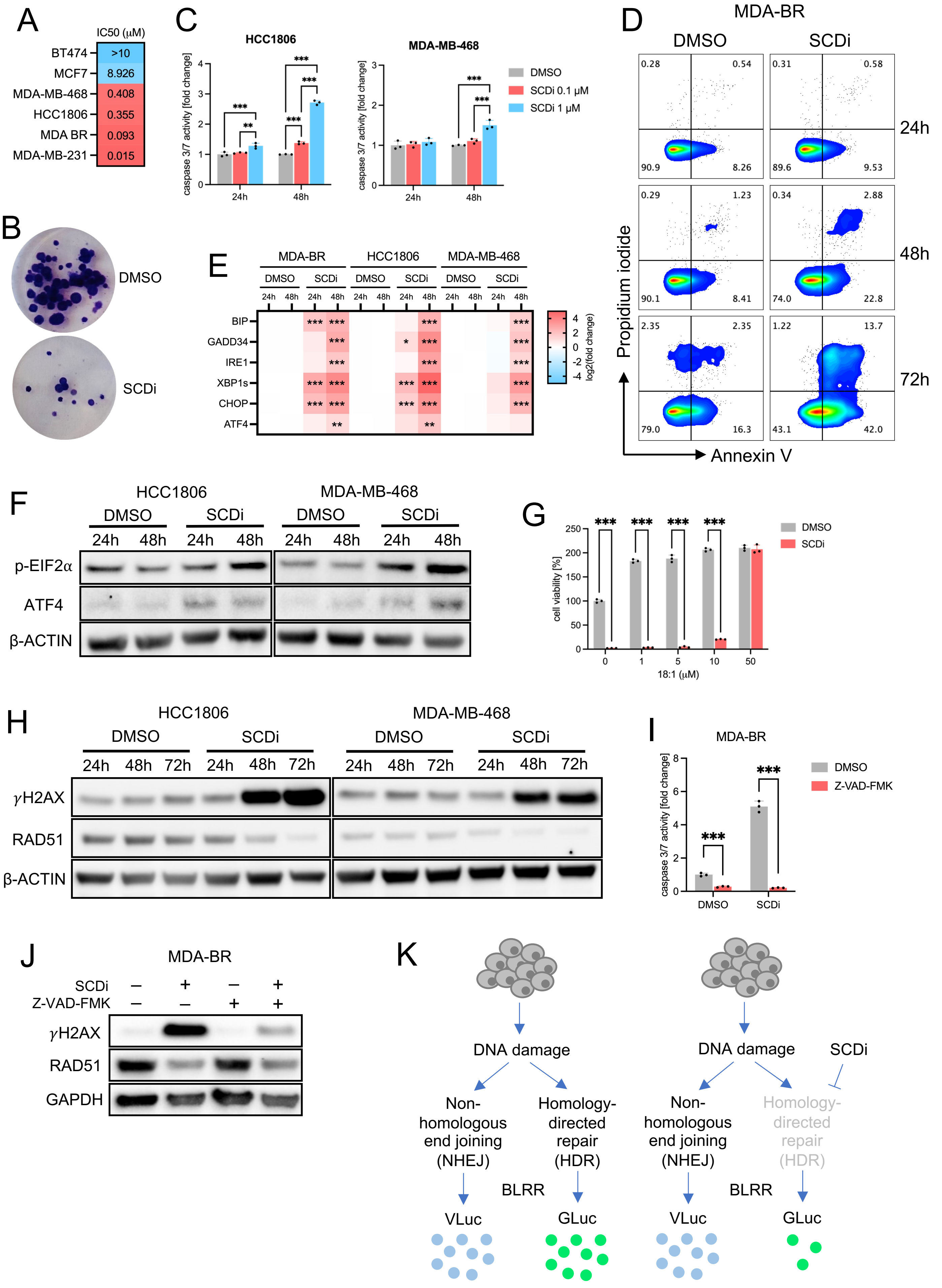

A
IC50 (μM)
BT474 >10
MCF7 8.926
MDA-MB-468 0.408
HCC1806 0.355
MDA BR 0.093
MDA-MB-231 0.015
B
DMSO
SCDi
C
HCC1806
MDA-MB-468
caspase 3/7 activity [fold change]
24h
48h
DMSO
SCDi 0.1 μM
SCDi 1 μM
D
MDA-BR
DMSO
SCDi
24h
48h
72h
Propidium iodide
Annexin V
E
MDA-BR
HCC1806
MDA-MB-468
BIP
GADD34
IRE1
XBP1s
CHOP
ATF4
log2(fold change)
F
p-EIF2α
ATF4
β-ACTIN
G
cell viability [%]
18:1 (μM)
H
γH2AX
RAD51
I
MDA-BR
Z-VAD-FMK
J
GAPDH
K
DNA damage
Non-homologous end joining (NHEJ)
Homology-directed repair (HDR)
BLRR
VLuc
GLuc
SCDi

**Figure S2. Pharmacological inhibition of SCD induces ER stress and is associated with DNA damage.**

(A) Half-maximal inhibitory concentration (IC50) of breast cancer cell lines treated with SCDi for 72h.

(B) Colony formation assay of MDA-BR cells treated with DMSO or SCDi (0.1 □M) for 72h.

(C) Caspase 3/7 assay of HCC1806 and MDA-MB-468 treated with SCDi. Mean □ SD (Two-way ANOVA, Tukey's test).

(D) Annexin V/Propidium iodide assay of MDA-BR cells treated with SCDi (1 □M).

(E) Relative gene expression of endoplasmic reticulum (ER) stress markers in MDA-BR, HCC1806, and MDA-MB-468 cells treated with DMSO or SCDi (1 □M), measured by qPCR.

(F) Western blot for ER stress markers in HCC1806 and MDA-MB-468 treated with SCDi (1□M).

(G) Cell viability assay of MDA-BR treated with oleic acid 18:1 with or without SCDi (1 □M). Mean □ SD (Student's t test).

(H) Western blot for DNA damage markers in HCC1806 and MDA-MB-468 cells treated with SCDi (1□M).

(I) Caspase 3/7 assay of MDA-BR treated with SCDi (1 □M) and/or with the pan-caspase inhibitor Z-VAD-FMK (50 □M). Mean □ SD (Student's t test).

(J) Western blot for DNA damage markers in MDA-BR cells treated with SCDi (1 □M) and/or with the pan-caspase inhibitor Z-VAD-FMK (50 □M).

(K) Schematic representation of the bioluminescent reporter (BLRR) used to track DNA damage repair dynamics. **, $p < 0.01$; ***, $p < 0.001$.

Figure S3

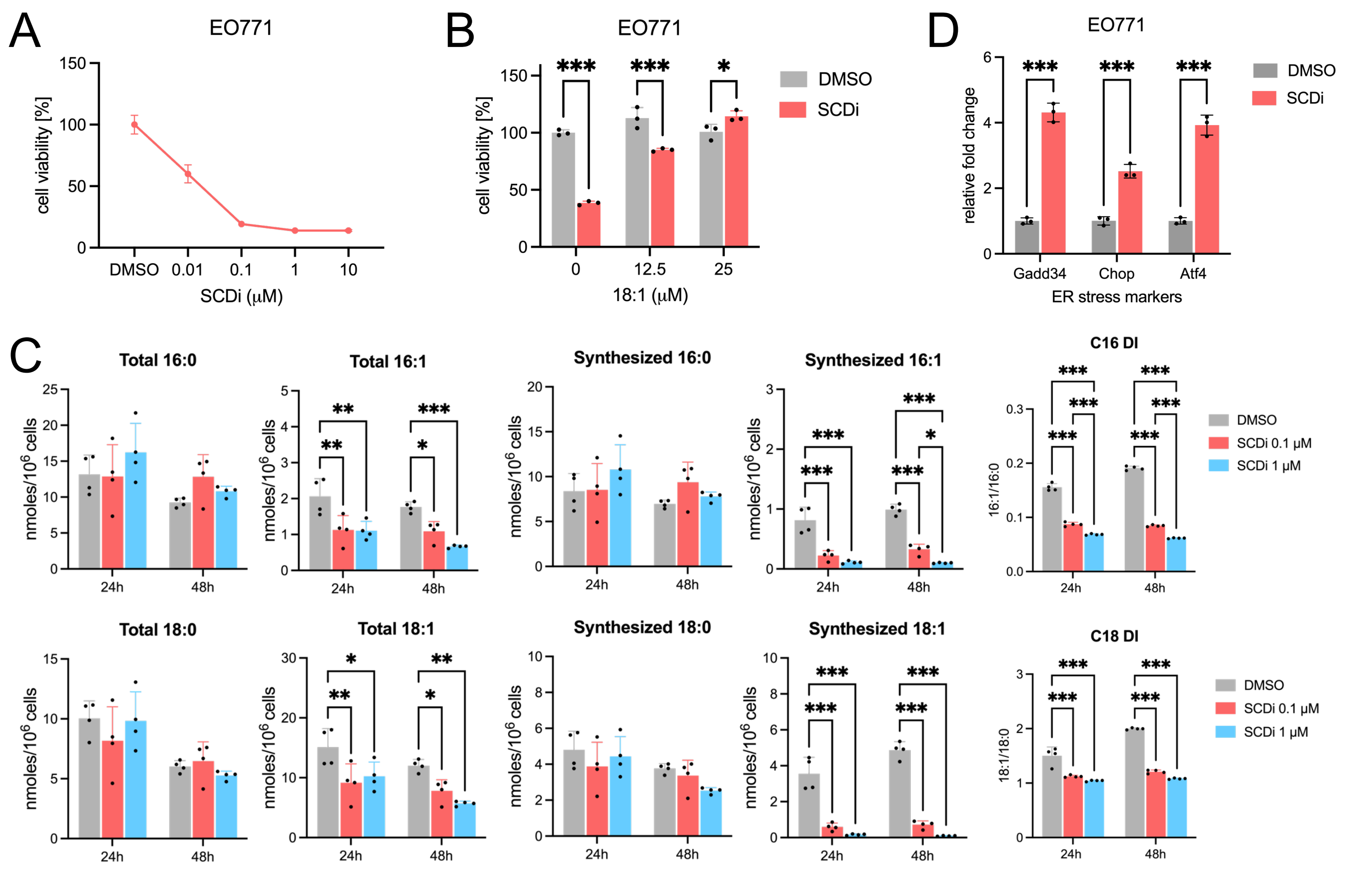


**Figure S3. Inhibition of SCD leads to cytotoxicity, decreases the desaturation index, and triggers ER stress in mouse mammary cancer cells.**

(A) Cell viability of EO771 treated with SCDi for 72h. Mean ± SD.

(B) Cell viability of EO771 treated with oleic acid 18:1 with or without SCDi (1 [illegible]M). Mean [illegible] SD (Student's t test).

(C) Gas chromatography/mass spectrometry of EO771 treated with SCDi for 24h and 48h, measuring total and synthesized palmitic acid (16:0), palmitoleic acid (16:1), stearic acid (18:0), and oleic acid (18:1). On the right, the desaturation index (ratio 16:1/16:0 and 18:1/18:0) is shown. Mean [illegible] SD (Two-way ANOVA, Tukey's test).

(D) Relative gene expression of endoplasmic reticulum (ER) stress markers in EO771 cells treated with DMSO or SCDi (1 [illegible]M) for 48h measured by qPCR. Mean [illegible] SD (Student's t test). *, $p < 0.05$; **, $p < 0.01$; ***, $p < 0.001$.

Figure S4

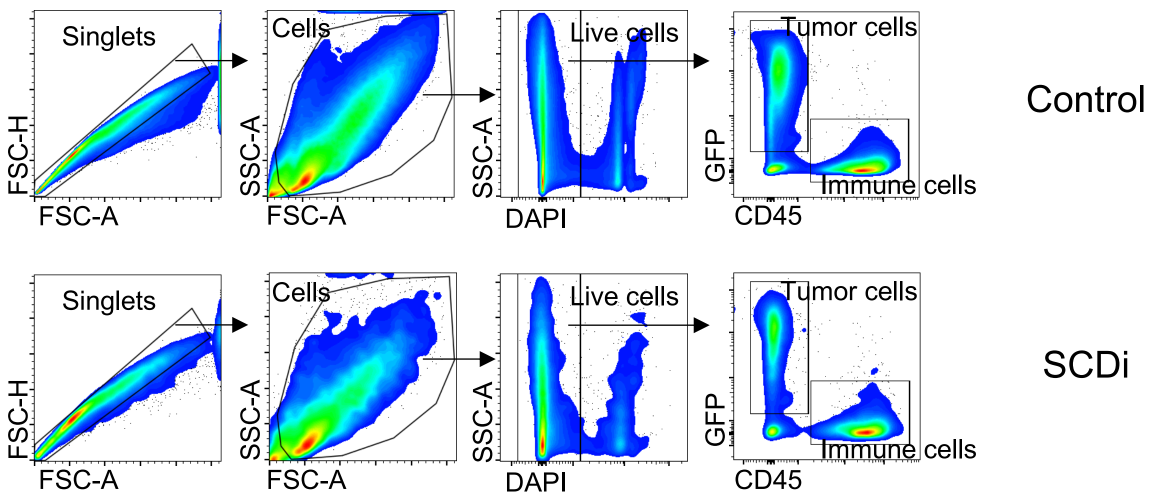


| Population | Control | SCDi |
|---|---|---|
| Immune cells | 55.6% | 61.2% |
| Tumor cells | 33.4% | 29% |

| Population | Control | SCDi |
|---|---|---|
| Immune cells | 2.9 x $10^6$ | 2.5 x $10^6$ |
| Tumor cells | 2.3 x $10^6$ | 1.2 x $10^6$ |

**Figure S4. Effect of SCD inhibition in a syngeneic mouse model of breast cancer brain metastases.**

Scatter plots representing the flow cytometry gating strategy used on brain tumors of mice intracranially injected with EO771-GFP either treated with vehicle (Control) or with SCDi for 14 days that were submitted for single-cell RNA sequencing analysis. Three mice per group were pooled before fluorescence activated cell sorting.

Figure S5

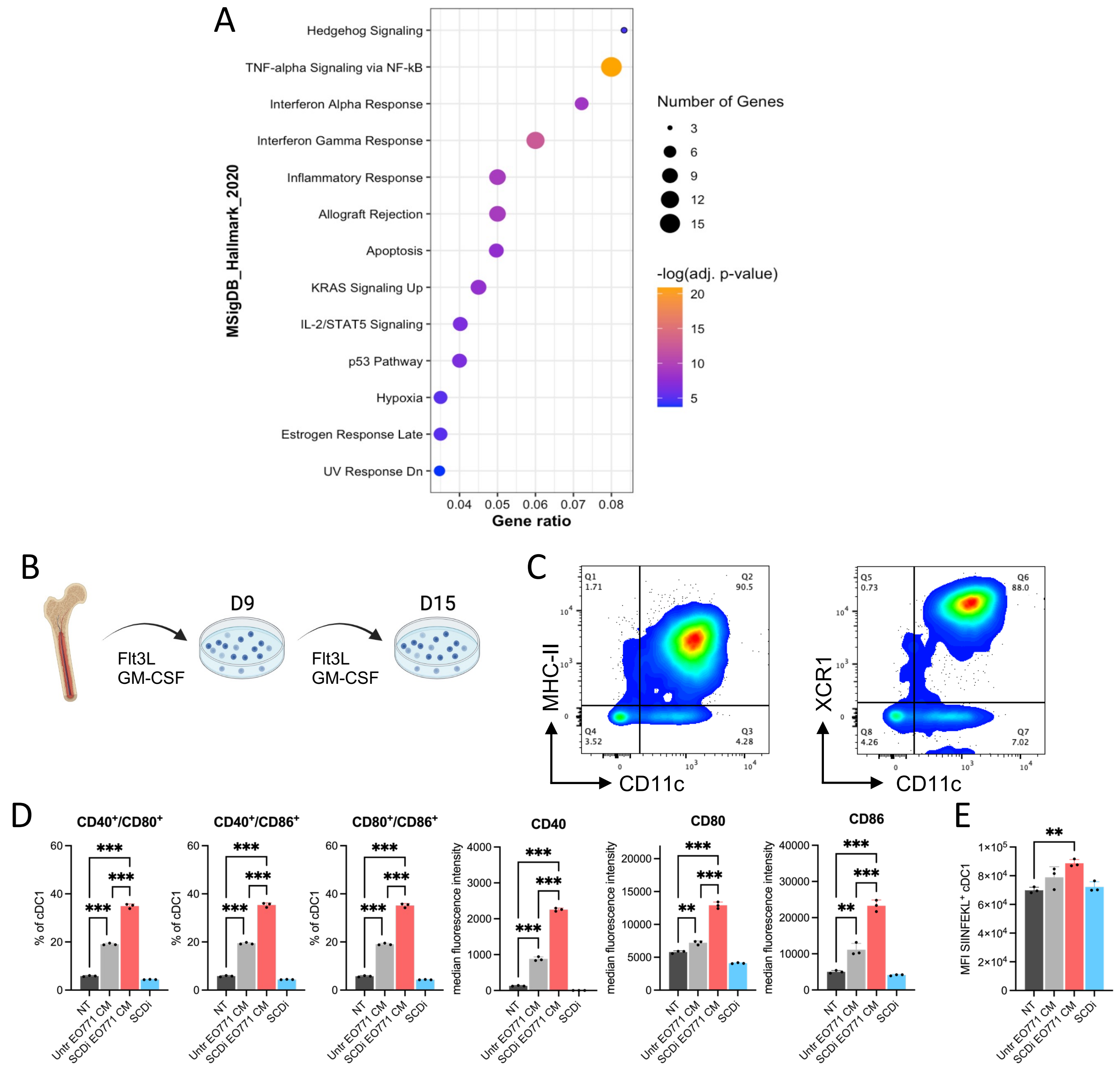


**Figure S5. Inhibition of SCD is associated with a higher dendritic cell activation.**

(A) Pathway analysis (MSigDB) of the differentially expressed genes (DEGs) in dendritic cells (SCDi versus control).

(B) Schematic depicting the differentiation protocol of bone marrow-derived dendritic cells (BMDCs).

(C) Scatter plots showing the expression of CD11c, MHC-II, and XCR1 on BMDCs at day 15 after differentiation, measured by flow cytometry.

(D-E) Histograms representing flow cytometry analysis of BMDCs that were either non-treated (NT), treated with DMSO-treated (Untr) EO771 conditioned medium (CM), with SCDi-treated EO771 CM, or with SCDi alone. Mean □ SD (Student's t test). **, $p < 0.01$; ***, $p < 0.001$.

### Figure S6

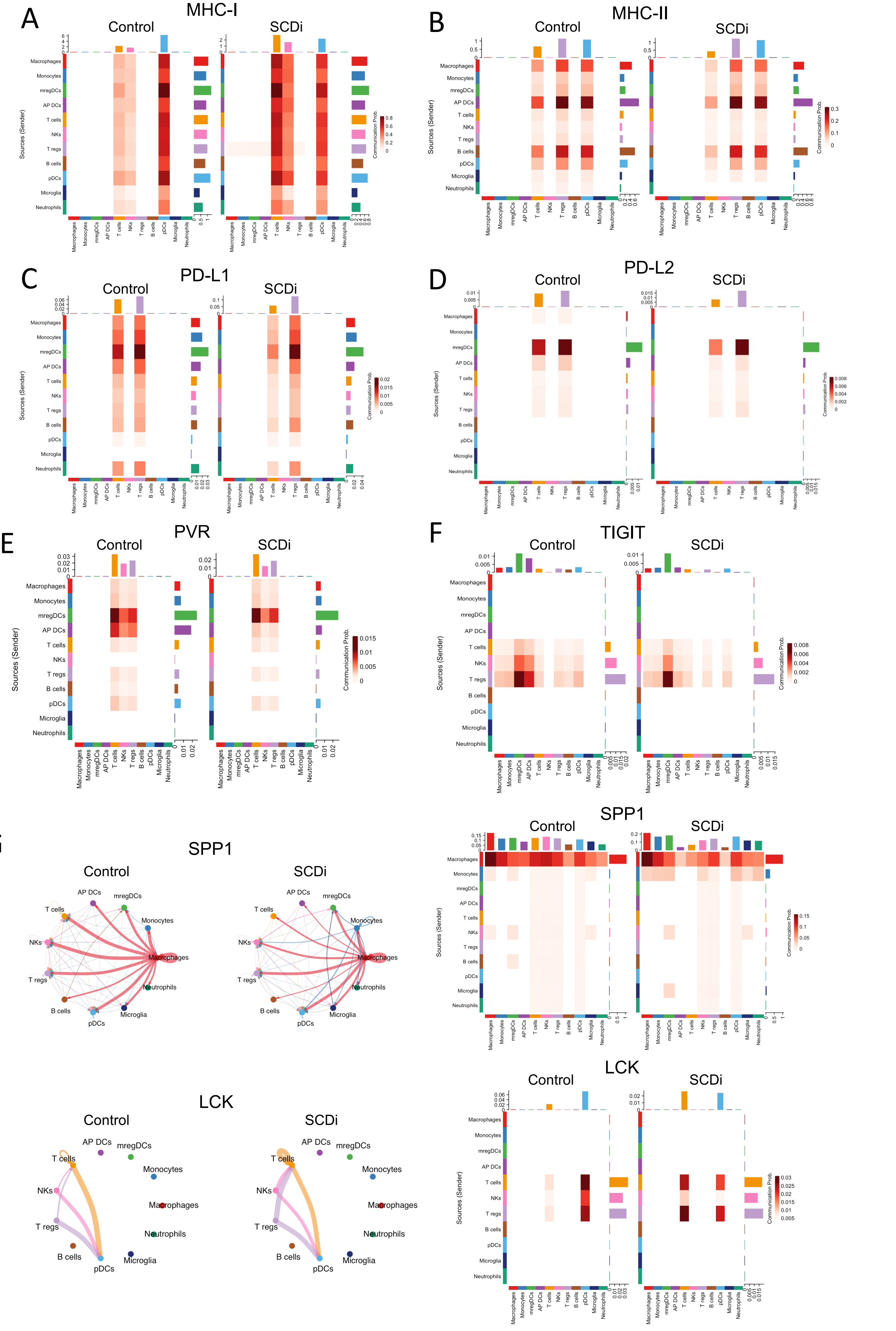

A
MHC-I
Control
SCDi
B
MHC-II
Control
SCDi
C
PD-L1
Control
SCDi
D
PD-L2
Control
SCDi
E
PVR
Control
SCDi
F
TIGIT
Control
SCDi
G
SPP1
Control
SCDi
H
LCK
Control
SCDi
Sources (Sender)
Communication Prob.
Macrophages
Monocytes
mregDCs
AP DCs
T cells
NKs
T regs
B cells
pDCs
Microglia
Neutrophils

**Figure S6. CellChat analysis between immune cell populations in the tumor microenvironment.**

(A-F) Heatmaps representing inferred communication probability between senders (Y axis) and receivers (X axis) via (A) MHC-I, (B) MHC-II, (C) PD-L1, (D) PD-L2, (E) PVR, and (F) TIGIT signaling pathways in control- and SCDi-treated tumors.

(G-H) Inferred (G) SPP1 and (H) LCK signaling networks among the cell population represented by the nodes on the circle plot (left). The thickness of lines connecting two cell populations represent the interaction strength (i.e., thicker lines represent stronger interactions). On the right, heatmaps representing inferred communication probability between senders (Y axis) and receivers (X axis) via (G) SPP1 and (H) LCK signaling pathways in control- and SCDi-treated tumors.

**Table S1**. Primer sequences used for qPCR analysis

| Gene | Species | Forward primer 5' | Reverse primer 5' |
|---|---|---|---|
| BIP | Homo sapiens | CATCACGCCGTCCTATGTCG | CGTCAAAGACCGTGTTCTCG |
| GADD34 | Homo sapiens | AGCCACGGAGGATAAAAGAACA | CTGAACGATACTCCCAGGACC |
| IRE1 | Homo sapiens | CATCCCCATGCCGAAGTTCA | CTGCTTCTCTCCGGTCAGGA |
| XBP1s | Homo sapiens | GGTCTGCTGAGTCCGCAGCAGG | GGGCTTGGTATATATGTGG |
| CHOP | Homo sapiens | GGAAACAGAGTGGTCATTCCC | CTGCTTGAGCCGTTCATTCTC |
| ATF4 | Homo sapiens | CCCTTCACCTTCTTACAACCTC | TGCCCAGCTCTAAACTAAAGGA |
| HPRT | Homo sapiens | GGCGAACCTCTCGGCTTT | AAGACGTTCAGTCCTGTCCA |
| ACTB | Homo sapiens | TGGCACCACACCTTCTACAA | CCAGAGGCGTACAGGGATAG |
| Gadd34 | Mus musculus | CTCTAAAAGCTCGGAAGGTACAC | GGCTTCGATCTCGTGCAAAC |
| Chop | Mus musculus | CCCATGCCCTTACCTATCGT | AGGTTTTTGATTCTTCCTCTTCGT |
| Atf4 | Mus musculus | GCTGCGGTAGGATCACG | ATTTCGTGAAGAGCGCCAT |
| Hprt | Mus musculus | TGATCAGTCAACGGGGGACA | TTCGAGAGGTCCTTTTCACCA |
| Actb | Mus musculus | CGCAGCCACTGTCGAGTC | GTCATCCATGGCGAACTGGT |
